# Optimized AAV capsids robustly transduce airway epithelial cells

**DOI:** 10.64898/2026.05.10.723853

**Authors:** Ashley L. Cooney, Yong Hong Chen, Brian C. Lewandowski, Shakayla Lamer, Griffin Boysen, Katarina Kulhankova, Amber Vu, Priyanka Newase, Patrick L. Sinn, Beverly L. Davidson, Paul B. McCray

**Affiliations:** University of Iowa, Stead Family Department of Pediatrics Iowa City, IA 52242, USA; Pappajohn Biomedical Institute, Iowa City, IA 52242, USA; Raymond G. Perelman Center for Cellular and Molecular Therapeutics, Children’s Hospital of Philadelphia, Philadelphia, PA 19104, USA; Department of Pathology and Laboratory Medicine, Perelman School of Medicine, University of Pennsylvania, Philadelphia, PA 19104, USA

**Keywords:** lung gene therapy, cystic fibrosis

## Abstract

Gene therapies have demonstrated transformative potential for a range of genetic disorders, including immunodeficiencies, hematopoietic conditions, and neuromuscular diseases. However, the application of these approaches to cystic fibrosis (CF) and other airway diseases remains constrained by the challenge of efficient gene delivery to target epithelial cells. Adeno-associated virus (AAV) vectors are widely used for in vivo gene delivery due to their favorable safety profile and capacity for long-term transgene expression in non-dividing cells. Nonetheless, current AAV capsids require high doses to achieve therapeutic efficacy in the airways, raising safety concerns. Here we report the development of novel AAV capsid variants with markedly enhanced transduction efficiency of airway epithelial cells. Using unbiased peptide-modified AAV libraries and round-over-round screening in well-differentiated primary cultures of human airway epithelia (HAE), we identified 20 novel capsids that efficiently transduced cells at doses 10- to 100-fold lower than those required by existing vectors (termed AAV-AE). These variants demonstrated high transgene expression in HAE, primary human basal cells, tracheal explants from nonhuman primates, and murine airways in vivo. These optimized AAV capsids represent a significant advancement in pulmonary gene therapy, offering a versatile platform for the delivery of gene addition and editing reagents to treat CF and other respiratory diseases.

## INTRODUCTION

Despite decades of effort, the use of gene therapy to treat inherited lung diseases, most notably cystic fibrosis (CF), remains difficult in large part due to the lack of efficient delivery vectors. CF is an autosomal recessive disease occurring in approximately 1 of every 3,200 live births in the US. Although CF affects multiple organ systems, the leading cause of morbidity and mortality is lung disease. CF is caused by mutations in the cystic fibrosis transmembrane conductance regulator (*CFTR*) gene, an anion channel that plays an important role in regulating the movement of salt and water across epithelial surfaces in the lungs, digestive system, and other organs. Disruption of CFTR function causes build-up of viscous mucus that obstructs airways, impairs host defenses, and leads to chronic bacterial infections and progressive tissue damage. Genetic therapies for CF directly address the impact of pathogenic variants of the *CFTR* gene. Such therapies include either gene addition (delivery of a functional *CFTR* cDNA or mRNA) or gene editing (repair of the endogenous chromosomal locus). While Trikafta and other small molecule CFTR modulators provide benefit to the majority of people with CF^1,2^, ∼10% carry mutations unresponsive to modulator therapies or are intolerant of the currently available options. Multiple ongoing gene therapy trials are testing the potential for restoring functional CFTR expression in the intrapulmonary airways, using lipid nanoparticles^3,4^, antisense oligonucleotides^5^, HSV-1 based vectors, lentiviral vectors^6^, and adeno-associated viral-(AAV) based vectors^7–9^. Despite these efforts, there is still significant need for improved efficiency in airway-tropic vectors to enable effective therapeutic delivery at lower vector dosages, improving safety, and reducing the cost of treatment.

AAV was the first approved viral vector treatment for a genetic disease^10^. AAV therapeutics have achieved success in numerous diseases including Leber congenital amaurosis, the hemophilias, sickle cell disease, β-thalassemia, spinal muscular atrophy, cerebral adrenoleukodystrophy, and others^11^. Five AAV-based CF clinical trials took place from 1999-2007, but ultimately the efficacy was too low and a therapy was not approved (reviewed in^12^). Advances including shortened promoters^13,14^, a *CFTR* mini gene lacking a portion of the R domain^15^, use of an augmenter to increase expression^16^, and improved AAV capsids^17,18^ have increased transduction and expression efficiencies in airway epithelia. Two AAV-based clinical trials for CF initiated by 4D Molecular Therapeutics (NCT05248230, clinicaltrials.gov) and Spirovant (NCT06526923, clinicaltrials.gov)^7,8^ are currently underway using evolved capsids based on AAVs 2, 5, or 6. However, even for these newer generation evolved capsids, airway epithelia continue to be a difficult tissue to transduce. The development of efficient delivery vectors is critical for the successful implementation of gene therapy to the lung and other tissues. Delivering therapeutic dosages at lower viral loads reduces the risks of off-target toxicity and immunogenicity, the primary barriers for translation of AAV gene therapies to clinic^19^.

In this study, we devised an unbiased approach to identify highly potent capsids capable of transducing human airway epithelial cells (HAE). These experiments build off methodology developed by our group that employs advanced AAV barcoding methods and round-over-round screening to identify novel capsids with enhanced tropism for target tissues^17,18^. In prior studies using these methods, we identified novel peptide-modified AAV (PM-AAV) capsids that transduced target CNS cell types at doses 1 to 2 logs lower than currently available capsids^17,18^. Here, we leveraged this PM-AAV capsid library approach to identify highly performant capsid variants targeting human airway epithelia. After three rounds of screening through primary HAE cultures using PM-AAV libraries based on 3 parental serotypes (AAV2, AAV6, and AAV9), we identified 20 novel AAV variants with remarkably enhanced tropism for airway epithelia (termed AAV-AE). We further validated the activity of these variants in human airway and human basal cell cultures, as well as lung tissue from non-human primates (NHPs) and mice. Several of these novel AAV-AEs transduced airway epithelia at levels higher than any previously delivered lung-tropic AAVs and at 10+ fold lower doses.

One novel variant in particular, AAV-AE-C5, exhibited remarkably enhanced tropism in human basal, secretory, and ciliated cells. Notably, this variant exhibited strong transduction even in the absence of the secondary transduction augmentor doxorubicin (Dox), which could simplify gene therapy treatments for lung diseases. When applying all available tools, including Dox and formulation in high NaCl^20^, AAV-AE-C5 transduced 3+-fold more HAE cells than parental serotypes and the engineered lung-tropic AAV2.5T^7,8^, and did so at 1/10^th^ of the dose. The significant improvements in lung-tropism exhibited by these novel capsids, most notably AAV-AE-C5, represent an important step towards overcoming the inefficiencies in therapeutic delivery to lung tissue that have hampered efforts to use gene therapy to treat hereditable pulmonary diseases.

## RESULTS

### Generating peptide display libraries

Using PM-AAV capsid libraries, we conducted a multi-round screen to identify novel capsid variants with enhanced tropism for HAE cell types. AAV capsids were modified by inserting a 7-mer peptide sequence into the variable region of loop VIII at positions 587, 587, and 588 in AAV2, AAV6, and AAV9, respectively. Peptide sequences were comprised of semi-random DNA bases “NNKNNKNNKNNKNNKNNKNNK” where “N” represents DNA bases A, C, T or G and “K” represents G or T. PM-AAV libraries for each serotype were pooled to create an input virus containing an estimated ∼6.8 million capsid variants^17,18^, which served as the initial (Round 1) input virus for multi-round PM-AAV screening in CF- and non-CF derived air liquid interface cultures of HAE (**Fig. 1A**). Viral DNA genomes recovered from HAE transduced with the Round 1 input virus were used to generate a Round 2 PM-AAV library containing 768,858 PM-AAV variants. PM-AAV variant performance in Round 2 was analyzed using next-generation Illumina amplicon sequencing (NGS) to measure the prevalence of variant genomes (DNA) and transgene expression (RNA). Based on analysis of these DNA and RNA sequencing results, 436 (AAV2=198, AAV6=142, AAV9=96) high performing PM-AAV capsid variants were identified and manufactured to serve as the input virus for a final round of screening. From NGS analysis of this final round of screening, 20 PM-AAVs (A1-6: AAV2, B1-6: AAV6, C1-8: AAV9) were selected for further validation (**Fig. 1B, C**). These candidates were chosen based on high transduction efficiencies across multiple HAE donors, with a qualitative preference for RNA data performance. Peptide diversity was also factored into the selection of these top 20 PM-AAV candidates, collectively termed AAV-AE (**Fig. 1B**).

**Figure 1:**
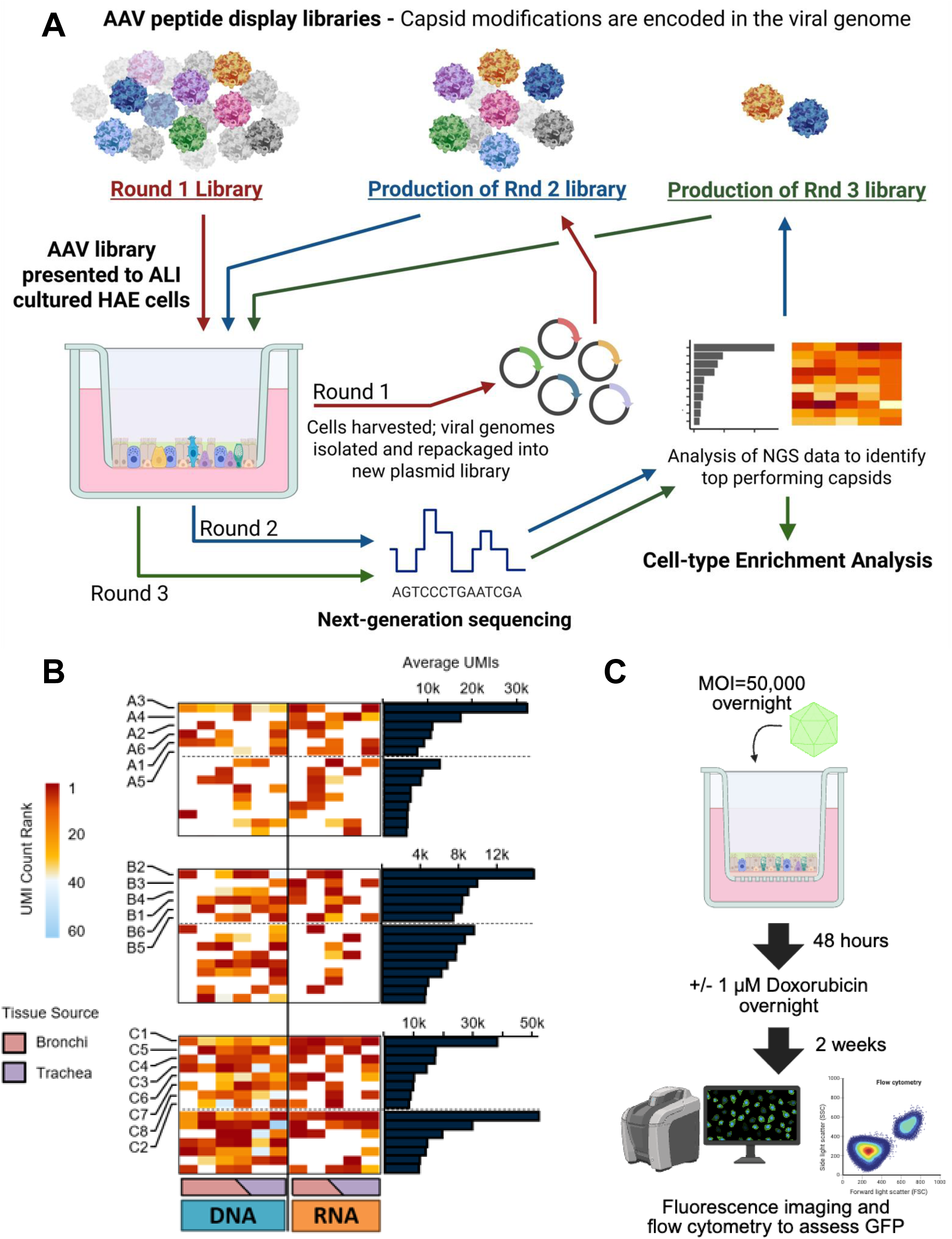
AAV capsid variant discovery pipeline. A) PM-AAV libraries for parental serotypes AAV2, AAV6, and AAV9 were generated by integrating a 7-mer peptide into the variable region of loop VIII. These PM-AAV libraries were combined to form a Round 1 input virus and applied to multiple HAE cultures from CF and non-CF donors. Intracellular AAVs were recovered, amplified, and manufactured into a Round 2 input virus for a second round of screening. NGS analysis of variant performance in Round 2 HAE samples identified top performing capsids which were used to generate input virus for a final (Round 3) round of screening. B) Capsid variant performance was assessed using NGS results from Round 3 HAE samples to identify top performing capsids for fluorescent validation. Heatmaps show AAV-AE capsid performance across HAE cultures grown from bronchial and/or tracheal tissue samples quantified using UMIs count rankings in each HAE dataset. Associated bar plots show the average UMI counts across all the HAE datasets shown in the heatmaps. Capsid variants selected for validation are labeled and plotted at the top of the heatmaps followed by the capsids not selected for validation with the highest AVG UMI counts. C) Schematic showing the standard protocol used to evaluate capsid variant transduction of HAE cultures. AAV-AE variants, parental serotypes, and AAV2.5T carrying GFP were tested using this standard protocol. Variations on this standard protocol including MOI, no Dox, or formulation with hypertonic saline were performed as described in the text.

### AAV variant screen on human airway epithelia compared to benchmark capsids

We benchmarked novel capsids to a previously established airway-tropic capsid, AAV2.5T^7,8^, and to the parental serotypes AAV2, AAV6, and AAV9. AAV capsids carrying GFP (MOI=50,000) were applied to the apical surface of HAE overnight. Gene expression was compared in the presence or absence of the proteosome inhibitor doxorubicin (Dox) applied 48 hours post-transduction to the basolateral media at 1 µM overnight. Dox is an established augmentor known to increase AAV transgene expression in airway epithelia^21^. Unless indicated otherwise, this is the standard protocol used for AAV transduction of HAE. Two weeks later, GFP expression was visualized by fluorescence microscopy and both total transduction and the transduction of individual HAE cell types was quantified by flow cytometry (**Fig. 2A-C, Supplemental Fig. 1**). These benchmarking experiments found that AAV2.5T transduced ∼27% of HAE cells, and the three parental serotypes transduced 4-8% of HAE cells, with basal, secretory, and ciliated cells transduced equivalently.

**Figure 2:**
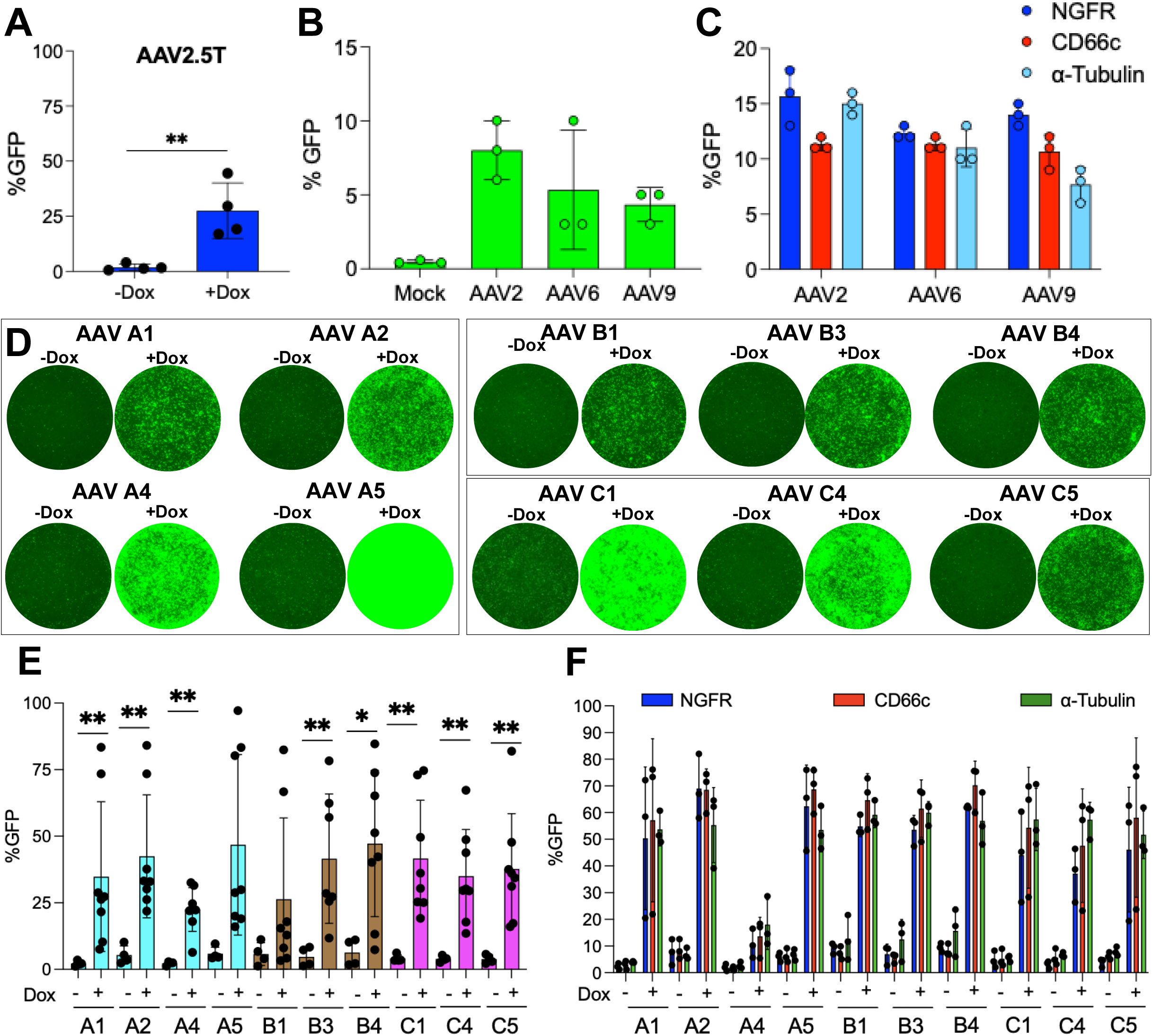
The top 10 AAV-AE variants demonstrate significantly improved HAE transduction efficiency compared to AAV2.5T and parental serotypes. Transduction of HAE cultures by parental and peptide-modified AAV variants packaged with GFP and transduced using the standard protocol (Fig. 1C) at an MOI of 50,000 either with or without Dox. Overall transduction was measured using GFP expression and quantified by flow cytometry. Transduction of individual airway epithelial cell types was quantified using flow cytometry of cells immunostained for relevant cell markers including basal cells (NGFR), secretory cells (Cd66c), and ciliated cells (⍺-tubulin). A) AAV2.5T-GFP transduced ∼27% of HAE cells in the +Dox condition, but very few cells in the -Dox condition (n=4, paired t-test, **p<0.005). B,C) Overall transduction (B) and cell type transduction (C) of HAE cultures by parental serotypes +Dox. Little transduction was observed in the -Dox condition (data not shown). D) Exemplar fluorescent images of HAE cultures for the top 10 AAV-AE variants -/+ Dox. Images were taken 2 weeks post-transduction. E,F) Quantification of overall transduction (E) and cell type transduction (F) -/+ Dox for AAV-AE variants A1, A2, A4, A5, B1, B3, B4, C1, C4 and C5. The increased transduction in the +Dox condition achieved significance for most of the capsids (E, n=8 per condition, one-way ANOVA, *p<0.05, **p<0.005).

The 20 identified AAV-AE capsids were applied to HAE using the standard protocol in the presence or absence of Dox (**Fig. 1C**, titers listed in **Supplemental Table 1**). Three weeks post-transduction, HAE were imaged for GFP expression (**Supplemental Fig. 2A-C**) and GFP+ cells quantified by flow cytometry. Benchmarked to AAV2.5T (**dotted line, Supplemental Fig. 2D**), we selected the top 10 capsid variants based on their GFP expression measured by flow cytometry (**Supplemental Fig. 2D**). These variants were A1, A2, A4, A5, B1, B3, B4, C1, C4, C5, with variants C4 and C5 notable for their expression levels in the absence of Dox treatment. Follow up experiments using HAE cultures derived from multiple donors and focused on these top 10 AAV-AE variants (**Fig. 2D,E**) replicated the results of the initial screening experiments, with the AAE-AE variants matching or out-performing AAV2.5T in the +Dox condition. Furthermore, unlike AAV2.5T and the parental serotypes, many of the AAV-AE variants also exhibited smaller, but consistent, levels of HAE transduction in the -Dox condition. When transduction of individual cell types important for airway function was measured, the majority of these AAV-AE variants had transduction levels exceeding 50% across all three major classes of airway cells: basal, secretory, and ciliated (**Fig. 2E**).

Inhaled hypertonic saline is a common supplementary treatment for CF that improves mucus clearance by rehydrating the airway surface layer. Our group has previously shown that hypertonic saline formulation also increases transduction of several viral vectors^20^. To test whether AAV-AE capsids may show similar improvements, we tested two of the top performing capsids from each serotype (A2, A4, B3, B4, C1, and C5) formulated with 4.5% NaCl (MOI=10,000, 2 hour apical transduction). The AAV-AE-C5 capsid showed a remarkable increase in gene transfer when formulated with 4.5% NaCl (**Supplemental Figure 3A, B**). To further characterize the effects of high NaCl formulation and Dox treatments on C5’s transduction efficiency, we decreased the dosage further (MOI=5,000) and measured C5 transduction efficiencies with and without 4.5% NaCl formulation in the presence of Dox. We observed significantly enhanced gene transfer when AAV-AE-C5 was formulated with 4.5% NaCl (**Fig 3A, B**). Even at this reduced MOI, C5 transduced ∼70% of HAE cells. We then tested the dose-response properties of 4.5% NaCl formulated AAV-AE-C5 +/- Dox at even lower concentrations (MOIs= 500, 1,000, 2,500, and 5,000 vg/cell). At 2 weeks, we quantified GFP expression and observed a dose dependent increase in GFP with or without Dox. Remarkably, at an MOI of 500, C5 achieved levels of transduction equivalent to the best of our benchmark capsids (AAV2.5T) tested at an MOI of 50,000 (**Fig 3C, D, Fig. 2A**).

**Figure 3:**
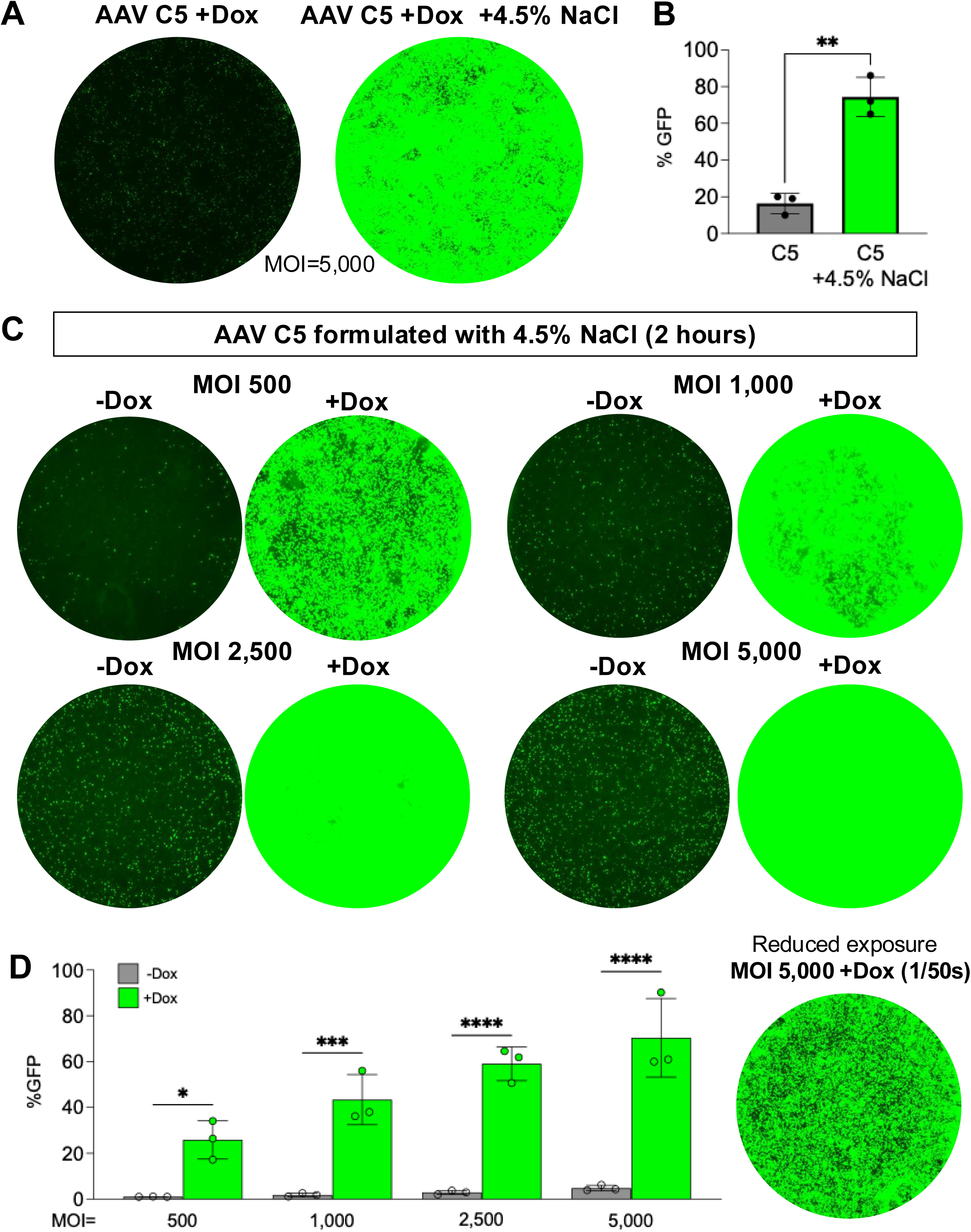
Formulating AAV C5 in hypertonic saline (4.5% NaCl) boosts gene transfer in HAE. AAV-AE-C5.GFP, formulated with or without 4.5% NaCl, was applied to the apical surface of HAE for 2 hours at an MOI of 5,000 and treated with 1 µM Dox 48 hours later. A) Representative fluorescent microscopy images of HAE cultures in the normal and hypertonic saline conditions. B) Transduction of HAE culture cells was quantified by measuring GFP expression using flow cytometry. Significantly more cells were transduced when AAV-AE-C5 was formulated in 4.5% NaCl (n=3, paired t-test, **p<0.005). C,D) The effect of 4.5% NaCl formulation on AAV-AE-C5.GFP transduction was further examined using the same methodology as above but including a -Dox condition and measuring dose-response properties by testing three lower MOIs: 500, 1000, 2500, in addition to the original 5,000. C) Representative GFP expression in HAE cultures visualized with fluorescent microscopy 2 weeks post transduction. D) Transduction efficiency measured 2 weeks post-transduction using GFP expression quantified by flow cytometry showed transduction in the 4.5% NaCl +Dox condition was significantly greater than in the -Dox condition at these MOIs (n=3, paired t-test, *p<0.05, ***p<0.0005, ****p<0.00005), and matched or exceeded results obtained for other top performing AAV-AE capsids in the +Dox condition (see Figure 2E), even at these 10+ fold lower MOIs.

### AAV C5 formulated with hypertonic saline expresses in HAE without doxorubicin

While the use of Dox has been approved in AAV-based clinical trials (NCT06526923), transducing airway epithelia without adjuncts would simplify vector approval. Here we tested the ability of our top performing AAV-AE variants to transduce HAE cells at a low dosage (MOI of 5,000) with and without Dox. Since doxorubicin leads to robust expression within days, we collected airway epithelial cells at 4 weeks for the +Dox condition (**Fig. 4A**). In the more challenging -Dox condition, we hypothesized that transfected cells may take longer to produce detectable quantities of GFP; so, GFP fluorescence for the -Dox group was monitored weekly by fluorescence microscopy and quantified at 8 weeks by flow cytometry (**Fig. 4B, C**). To ensure high levels of GFP did not cause cytotoxicity, we performed an LDH release assay of all AAV capsids at 4 weeks during the 8-week time course and found that no capsids produced elevated LDH release (**Supplemental Fig. 4**). In all capsids, we observed a gradual increase in GFP expression over the 8-week time course. In the -Dox condition, AAV-AE-C5 had the highest transduction percentage, which was further improved by formulation in 4.5% NaCl. As previously observed (**Supplemental Fig. 3**), in the +Dox condition some of the A series capsids had higher transduction than C5 in standard formulation. However, C5 formulated with 4.5% NaCl had the highest average transduction percentage, achieving significance in nearly all comparisons across both conditions (one-way ANOVA, **Fig. 4B, C**). Notably, AAV-AE-C5 formulated in 4.5% NaCl without Dox may reach sufficient transduction levels for an efficacious CF gene therapy application.

**Figure 4:**
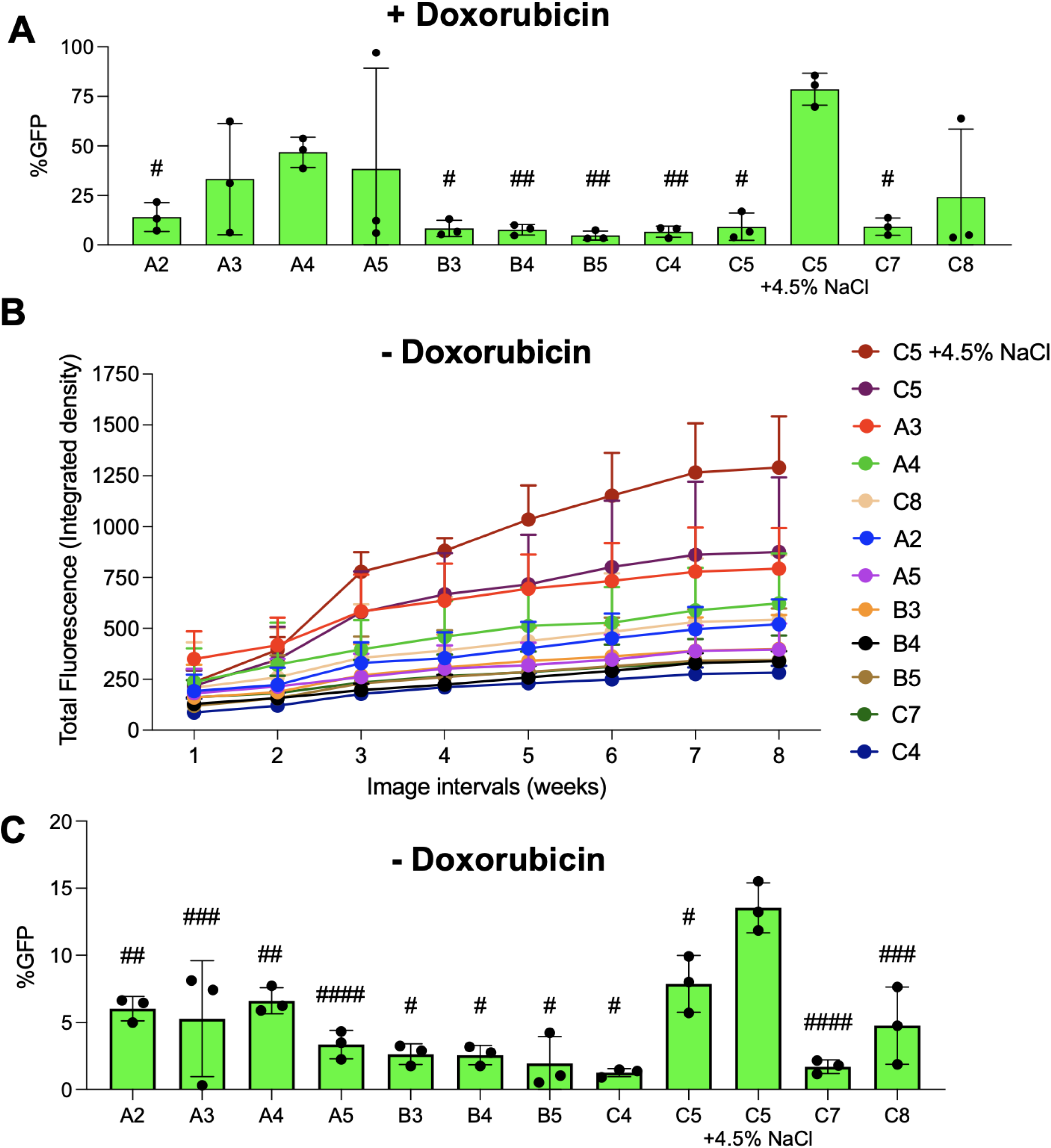
HAE culture transduction by AAV-AE capsids at 10-fold reduced dosages (MOI 5,000) with and without doxorubicin. Primary human airway epithelia were transduced apically with a selection of GFP-packaged AAV-AE capsids at MOI=5,000 overnight (for isotonic formulated capsids) or for 2 hours (for AAV-AE-C5.GFP formulated in 4.5% NaCl). Cultures were left untreated for the no doxorubicin condition (-Dox) or treated with 1 μM Dox overnight 48 hours post-transduction (+Dox). At 4 weeks for +Dox cultures (A), or 8 weeks for -Dox cultures (C), GFP expression was quantified by flow cytometry, n=3. B) To better visualize the time course of expression in the more challenging -Dox condition, airway cultures were imaged weekly for 8 weeks and mean fluorescence intensity was measured from each image and quantified using integrated density (Total Fluorescence). For many of the AAV-AE capsids, GFP expression continued to increase past the 4-week time point. In both the +Dox condition (A) and the -Dox condition (B), AAV-AE-C5 formulated in 4.5% NaCl had significantly higher transduction than most (+Dox) or all (-Dox) of the other AAV-AE’s tested (one-way ANOVA with Tukey’s multiple comparisons test, *p<0.05, **p<0.005, ***p<0.0005, ****p<0.00005).

### Capsid tropism in human airway cell cultures

To further characterize the cell types transduced by some of our top performing AAV-AE variants, we used confocal microscopy to identify GFP expressing cell types for a representative capsid from each AAV-AE group (A5, B3, C5, and C5+4.5% NaCl) and AAV2.5T for comparison. At a high magnification and using phalloidin staining to delineate localized F-actin structures, we observed quantities of GFP expressing cells consistent with the percentage of GFP detected by flow cytometry (**Fig. 5A**). Additionally, we observed ciliated (red arrows), secretory (orange arrows), and basal cells (blue arrows) transduced as shown on the XZ plane (**Fig. 5B**).

**Figure 5:**
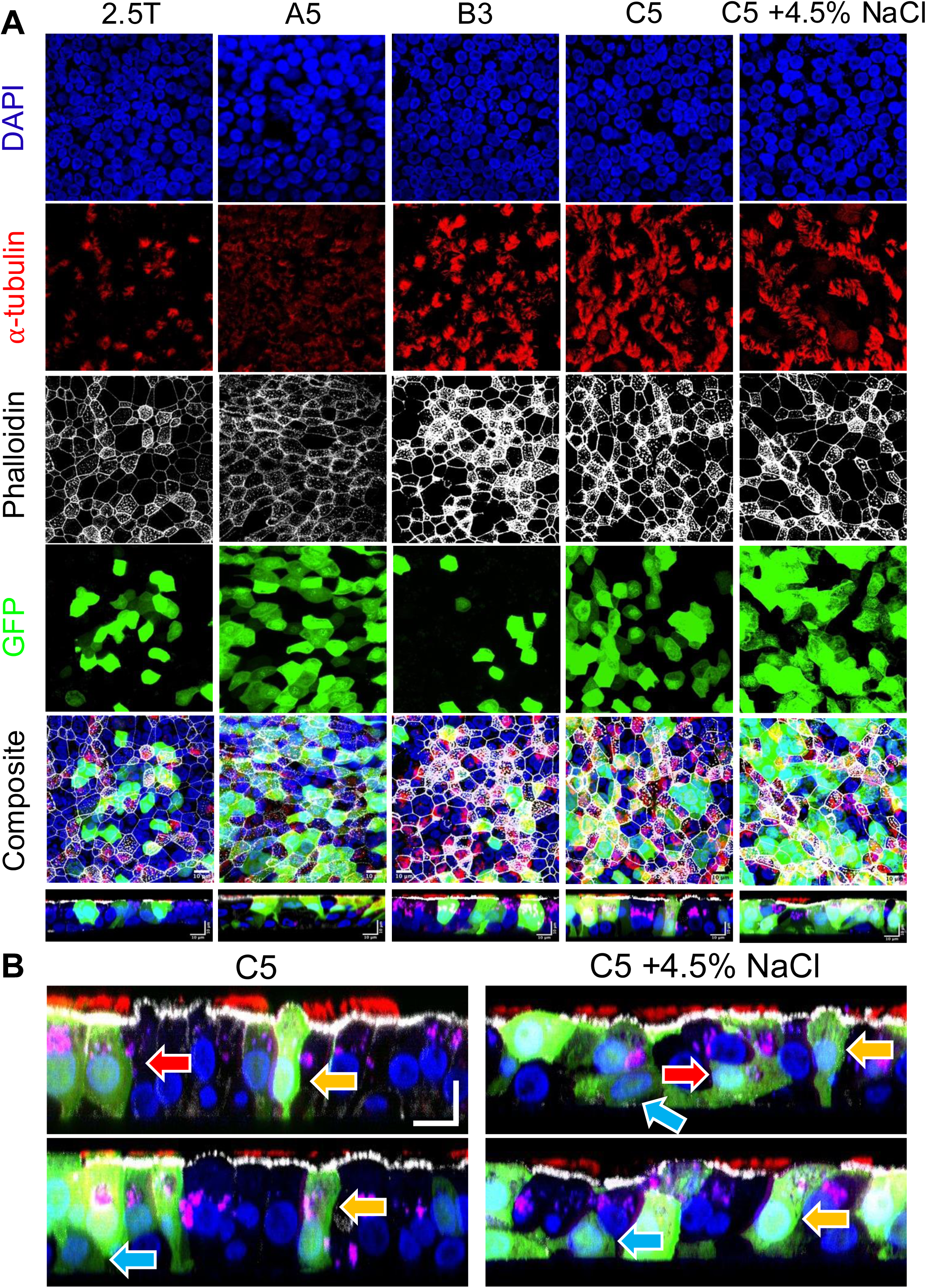
HAE morphology following AAV transduction. GFP-packaged AAV capsids including AAV2.5T, A5, B3, C5, and C5 +4.5% NaCl were used to transduce HAE cultures at MOI=50,000 and treated with 1 μM Dox 48 hours later. A) 2 weeks post-transduction, cells were fixed and immunostained with α-tubulin (red) and F-actin/phalloidin (white), mounted with DAPI (blue) and viral GFP transgene expression was visualized with fluorescent microscopy. Bottom panel is XZ plane from composite image. B) XZ plane from C5 and C5+4.5% NaCl show GFP+ ciliated cells (red arrows), secretory cells (orange arrows), and basal cells (blue arrows). Scale bars = 10 μm.

Basal cells are the progenitor cells of the conducting airway epithelium^22^. These airway stem cells are important targets for gene therapy because they can ensure that therapeutic efficacy continues even after turnover of the terminally differentiated surface airway epithelia. Given the importance of transducing this target population, we screened all 20 AAV-AE capsids on primary airway basal cells in submersion culture. AAVs were applied at MOI=10,000 overnight with a 1 µM Dox treatment. The highest overall levels of transduction (75%+) were exhibited by B group capsids B2, B5, and B6. Outside of those standouts, however, C5 transduced the largest percentage of basal cells (78%), though C8 and all the A group capsids had transduction rates of 50%+ (**Fig. 6A-B**). In total, 11 of the 20 capsids demonstrated basal cell transduction levels of 50% or greater.

**Figure 6:**
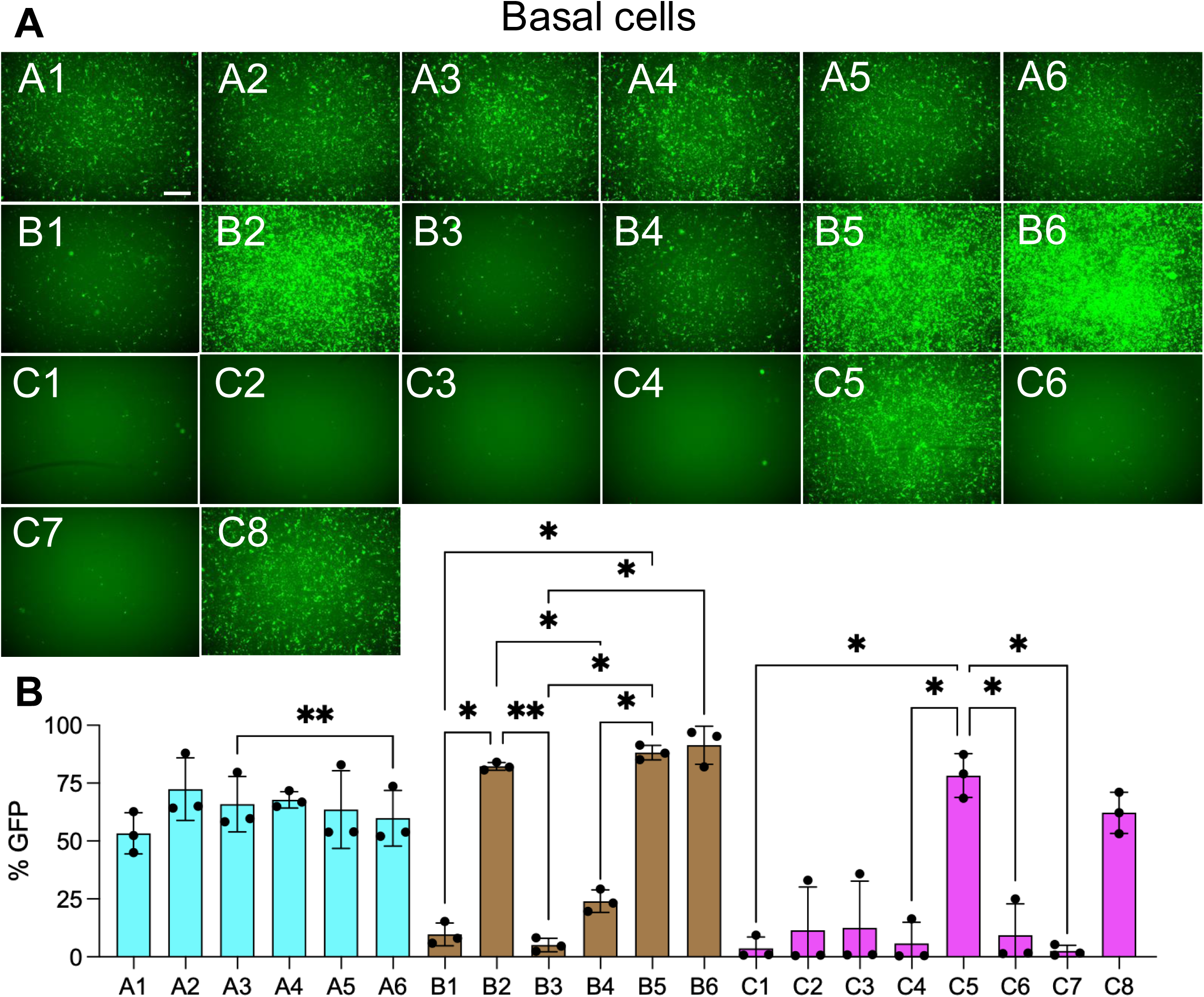
AAV-AE capsid transduction of human airway basal cells. Primary human airway basal cells were seeded on plastic and allowed to adhere overnight. The day after seeding, cells were transduced with AAV-AE variants (MOI=10,000), treated with 1 µM Dox and imaged 4 days post-transduction. A) Representative images of GFP fluorescence from each of the 20 AAV-AE capsids taken 4 days post-transduction (scale bar=500 µm). B) Comparison of basal cell culture transduction for the AAV-AE capsids (n=3 per capsid) measured using GFP fluorescence quantified by flow cytometry. One-way ANOVA statistical comparisons are within each capsid group A, B, or C, *p<0.05, **p<0.005.

### Species tropism and in vivo transduction efficiencies

Species dependent variation in cell tropism may limit translational studies in animal models. To determine if our AAV-AE capsids would also work in animal models, we first tested five of the highest performing capsids from HAE and basal cell cultures (A3, A4, A5, C5, and C8) on non-human primate tracheal explants. These capsids were formulated in 5% NaCl and transduction was assessed with and without Dox (**Figure 7A, B**). All the variants tested successfully transduced NHP airway cells, with a general trend towards improved transduction in the +Dox condition, but still notable levels of transduction in the absence of Dox.

**Figure 7.**
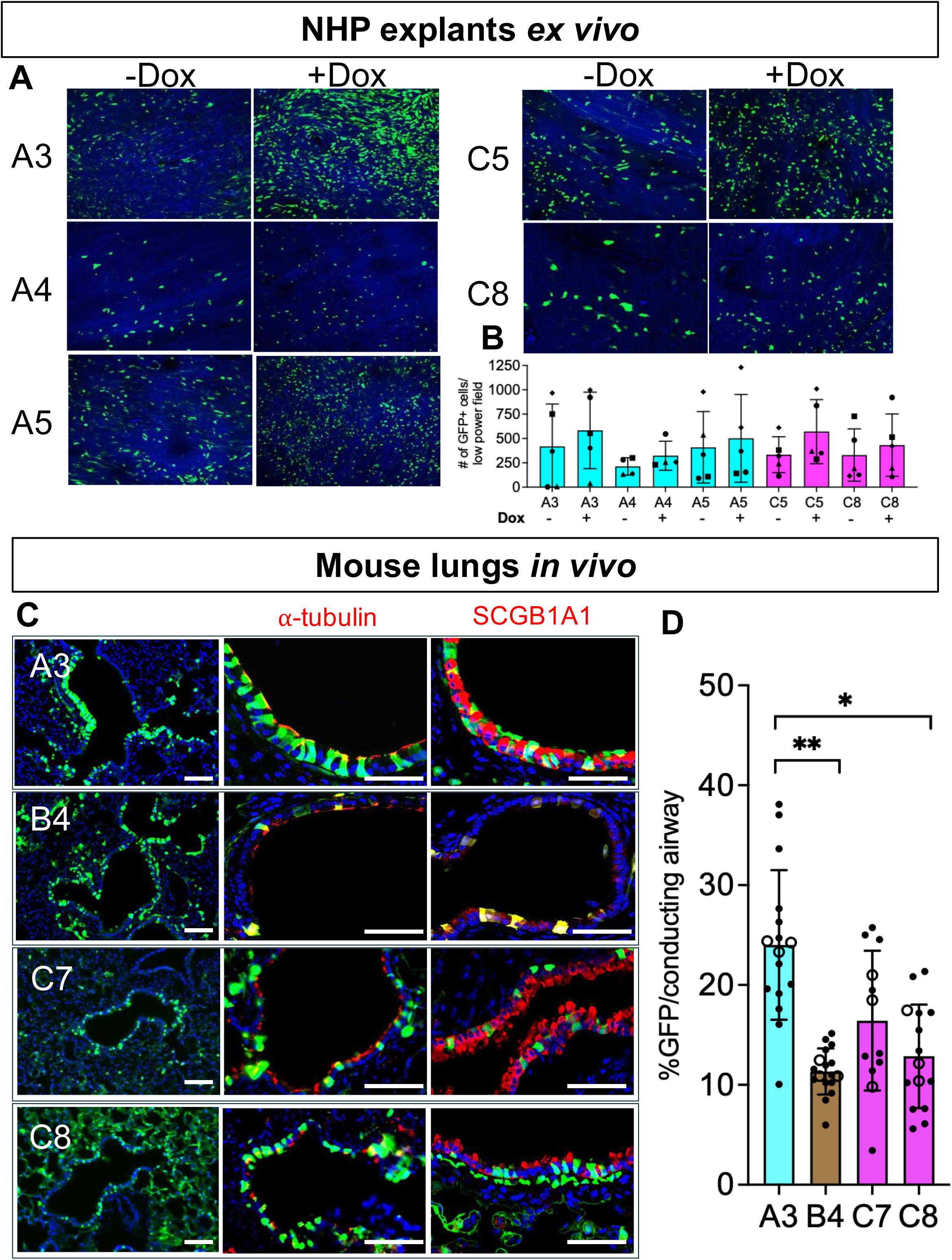
Non-human primate and mouse airway epithelia are transduced by AAV capsids. Non-human primate (NHP) tracheal explants (n=3) were cultured on Surgifoam^41^ and transduced with leading human AAV-AE capsids. NHP explants were transduced with 2x10^10^ vg (A3, A4, A5, C5, or C8) formulated with 5% NaCl and were compared with or without Dox (1 μM). Two weeks post-transduction, explants were immunostained with α-tubulin, mounted with DAPI, and imaged. A) Representative images showing GFP expression in NHP tracheal explants for each capsid + and -Dox. B) Viral transduction was quantified by counting the number of GFP positive cells within N representative X-by-X visual fields (n=5 explants per capsid). All capsids successfully transduced NHP cells, with the highest overall levels being observed in the +Dox condition. C,D) AAV variants formulated in 6% NaCl were instilled intratracheally into mouse airways at 1x10^11^ vg without Dox (n=3). Individual dots represent each airway counted, open circles indicate average per mouse. C) Representative images of mouse airway tissue for 4 of the more successful AAV-AE capsids in murine tissue (A3, B4, C7, C8). Blue=DAPI, green=GFP, red=tubulin (ciliated cells, middle panels) or SCGB1A1 (secretory cells, right panels). Scale bars =200 μm. F) GFP expression in conducting airways quantified using positive GFP cell counting in Fiji (one-way ANOVA; *p<0.05, ***p<0.0005, ****p<0.0005).

To test AAV-AE capsid variant transduction efficiency in murine airways, we selected 13 vectors with titers high enough to reach 1x10^11^ vg in less than a 100 µl volume (A3, A4, B3, B4, B5, C1-8, titers listed in **Supplemental Table 1**). Each vector was formulated with 6% NaCl and delivered via intratracheal intubation^20^ without Dox. Two weeks after vector administration, mice were euthanized and lungs collected, fixed, embedded, and sectioned for immunostaining. We observed the highest levels of transduction in the conducting airways for the A3, B4, C7, and C8 variants, including transduction of ciliated, secretory, and basal cells (**Fig 7C**). Little-to-no transduction was observed for capsids A4, B3, B5, and C3 while capsids C1, C2, C4, C5, and C6 transduced cells outside the conducting airways (data not shown). We quantified GFP expression in the conducting airways for capsids showing the greatest transduction efficiencies. Four of the capsid variants (A3, B4, C7, C8) transduced ∼10-40% of the airway epithelia, with A3 exhibiting the highest overall transduction efficiency (**Fig 7D**).

## DISCUSSION

Early efforts to treat genetic disorders in lung tissue were hampered by inefficiencies in both the therapeutics and delivery vectors. While significant advances have been made in therapeutic cassettes for heritable lung diseases such as CF^13–16^, effective delivery of these payloads to the lungs remains a challenge. AAVs are the most used viral vector for gene therapy delivery both to the lung and other tissues. The first AAV-CFTR clinical trials tested the AAV2 capsid^23–25^ and demonstrated low efficacy. Since then, transduction of conducting airway tissue by several additional AAV capsid variants has been studied including AAV1^26^, AAV5^27^, AAV6^28^, AAV9^29^ and others^7,9,30^. Attempts have been made to improve on the poor lung-tropism of these native AAV serotypes, including the use of evolved capsids^31^ and point mutations^32,33^, but these AAV-engineering efforts have only yielded modest improvements in airway transduction. The inefficiency of currently available lung-tropic AAVs means that high doses are necessary to achieve therapeutically sufficient transduction levels; this both increases the risk of off-target toxicity and the cost of treatment.

In this study we used an unbiased PM-AAV library screening approach^17,18^ to identify novel capsids with improved tropism for airway epithelia. After multiple rounds of screening in HAE cultures, we identified 20 AAV-AE capsids with significantly enhanced lung-tropism. We screened these capsids in multiple systems, including HAE, human basal cells, NHP tracheal explants, and mouse lungs. We identified several capsids with 10- to 100-fold improved tropism over the parental AAV2, 6, and 9 capsids, as well as AAV2.5T, a current AAV candidate for CF gene therapy (NCT06526923). One capsid, AAV-AE-C5, was particularly noteworthy. At 1/100^th^ the dose, C5 matches the transduction efficiency of the engineered, lung-tropic AAV2.5T capsid, and at 1/10^th^ the dose C5 exceeds AAV2.5T’s transduction efficiency by more than 2-fold. C5 also strongly transduces human basal cells, which may be necessary for long lasting clinical improvements, and is effective in NHP airway epithelial cells.

One strategy that has been employed to overcome the difficulties in targeting lung tissue with AAVs is the use of augmentors, specifically the proteosome inhibitor doxorubicin (Dox). Proteosome inhibitors have been shown to increase AAV-mediated transduction through multiple mechanisms, including the inhibition of capsid ubiquitination and the facilitation of microtubule transport of capsids to the nuclear envelope^21,34,35^. Our results replicate previous findings demonstrating the significant improvement in AAV-AE transduction of airway epithelia with Dox treatment^7^. In the absence of Dox, the parental serotypes AAV2, AAV6, and AAV9 and AAV2.5T transduce almost no airway epithelial cells. While Dox has been approved as an adjunct in a clinical trial^8^, administration of gene therapy would be simpler (and potentially safer) without the use of an augmenter. Importantly, we show that some of our AAV-AE capsids can transduce HAE in the absence of Dox. Once again, our C5 capsid was particularly impressive; at 1/10^th^ the dose at which parental and AAV2.5T capsids exhibited no transduction without Dox, AAV-C5 reached ∼15% HAE transduction.

Our group has also explored other strategies for improving AAV transduction of airway tissue without the use of adjuncts. We recognized that the high NaCl solutions commonly used in CF treatment also had the potential to facilitate AAV transduction^20^. We tested this strategy on some of our top performing capsids by formulating them in high NaCl. While most of the capsids we tested showed very modest improvements in HAE transduction when formulated in 4.5% NaCl, AAV-AE-C5 exhibited a dramatic 4+-fold increase. These results highlight the importance of considering vector formulation as an additional strategy for improving tissue transduction. This study has limitations. We did not identify a mechanism of entry or investigate the dependence of these capsids on AAVR for entry^36^. It is possible that the displayed 7mer peptide binds co-factors which enhance receptor targeting or entry to airway epithelia, however this will be investigated in future studies. Second, all cell types transduced in mouse airways by each capsid type were not characterized. Several capsids, specifically the C group, exhibited parenchymal transgene expression that likely represents alveolar epithelial cell transduction, an important target for surfactant dysfunction diseases such as ABCA3 deficiency, surfactant protein B deficiency, and surfactant protein C deficiency. Understanding capsid tropism will help identify appropriate capsids for mouse studies in disease models.

AAV capsids with remarkably enhanced transduction efficiency in airway epithelia could alleviate toxicity observed with high doses of AAV and improve the therapeutic index for lung applications^37^. Here we identified several PM-AAV capsids with robust transduction efficiencies in well-differentiated primary human airway epithelia, primary human basal cells, NHP tracheal explants, and mouse lungs. In summary, implementing these AAVs for gene transfer to the lung could advance several therapies for genetic lung diseases, including CF.

## MATERIALS AND METHODS

### Ethics Statement

Primary airway epithelia from human CF and non-CF donors were isolated from discarded tissue, autopsy, or surgical specimens. Cells were provided by The University of Iowa *In Vitro* Models and Cell Culture Core Repository. Information that could be used to identify a subject was not provided. All studies involving human subjects received University of Iowa Institutional Review Board approval (Protocol #230167). Mice experimental protocols were reviewed and approved by the University of Iowa Institutional Animal Care and Use Committee (IACUC), in accordance with the United States Department of Agriculture and National Institutes of Health guidelines.

### Amplicon sequencing data processing and analysis

Amplicon libraries were sequenced on a Novaseq 6000 using a 300-cycle reagent kit and 150bp paired-end read chemistry. Target read depth was 10 and 5 million reads per HAE sample for Rounds 2 and 3, respectively. A customized pipeline written in Python / R was used to analyze amplicon sequencing data. Initial quality control was performed by filtering Illumina FASTQs to remove low-quality reads by testing for perfect string matches against serotype specific constant regions of the amplicons. Nucleic acid sequences encoding the capsid peptide-modifications were parsed from passing reads and their frequencies tabulated. Capsid variant performance was assessed using two primary metrics: abundance and enrichment. Abundance (UMI counts) was quantified as the UMI-collapsed read count for each unique nucleic acid sequence. Enrichment represents the fold change in a capsid’s relative abundance in an HAE sample vs the input vector.

The Round 3 input virus was created by analyzing NGS results for each Round 2 HAE sample and selecting the top PM-AAVs based on the abundance metric in DNA data and in RNA data and then doing the same for the enrichment metric. In addition, PM-AAV performance across HAE samples was assessed using metric averages, and top performing capsids added to the Round 3 input virus. Selection of capsid variants for fluorescent validation involved a qualitative and quantitative assessment of Round 3 screening NGS results. The goal was to identify candidates that exhibited a balance of strong performance in DNA and/or RNA data, included top performers for all three parental serotypes, and represented a diverse set of peptide sequences. Because the goal of these screens is the delivery of therapeutic transgenes, added weight was given to RNA data performance when making these selections.

### Human airway epithelial cells

The University of Iowa *In Vitro* Models and Cell Culture Core cultured and maintained human airway epithelial cells (HAE) as previously described^38^. Briefly, following enzymatic disassociation of trachea and bronchus epithelia, the cells were seeded onto collagen-coated, polycarbonate Transwell inserts (0.4 μm pore size; surface area = 0.33 cm^2^; Corning Costar, Cambridge, MA). HAE were submerged in Ultroser G (USG) medium for 24 hours (37°C and 5% CO_2_) at which point the apical media is removed to encourage polarization and differentiation at an air-liquid interface. Transepithelial electrical resistance was measured using an Ohmmeter (Ω·μm^2^).

### Barcoding methods

To create the original library, random heptamer sequences were inserted into loop VIII in airway-tropic parental AAV capsid plasmids. These plasmids, along with helper plasmids were used to generate the Round 1 AAV library. Approximately 2.5M variants/serotype were generated and aliquots were applied to non-CF- and CF-derived primary HAE cultures. Intracellular virus was recovered from the HAEs, amplified using primers capable of detecting all serotypes, and the resulting viral genomes repackaged into AAVs for another round of screening. The resultant round 2 library contained approximately 750,000 variants and was again applied to a mixture of CF and non-CF HAEs. Transduced cells were harvested, viral DNA and RNA recovered as before, and the input from this round, as well as the output, was sequenced. From this output, 436 sequences total, representing the top-enriched variants for each serotype on HAE, were identified and used to generate input virus for a final round of screening.

### Viral vector production and transduction of HAE

AAV vectors were produced by the Children’s Hospital of Philadelphia Vector Core or the University of Iowa Viral Vector Core (https://medicine.uiowa.edu/vectorcore/). Vectors were titered by ddPCR to quantify vector genomes (vg)/ml. Each vector was applied to the apical surface of HAE airway epithelia (MOIs=500-50,000) either overnight or for 2 hours if formulated in 4.5% NaCl. 48 hours later, cells were left untreated or treated with 1 µM Doxorubicin (Dox) (D3447, Millipore Sigma) overnight. Primary human airway basal cells were transduced with MOI=10,000 and treated with 1 µM Dox. GFP expression was visualized and quantified 2-3 weeks post-transduction. To test AAVs formulated in 4.5% NaCl, vectors were prepared in 50 μl volumes composed of 9 μl of 25% NaCl, a volume of vector based on target MOI, and DMEM media. This mixture was applied to the apical surface of HAE for 2 hours and an overnight 1 µM Dox basolateral treatment was done 48 hours post-transduction.

### Fluorescence microscopy and flow cytometry

GFP images were acquired using a Keyence All-in-one Fluorescence Microscope BZ-X series (Osaka, Japan). 0.33 cm^2^ transwells were imaged at 2X magnification. GFP expression was quantified by flow cytometry as previously reported^39,40^. Briefly, cells were stained with a fixable LIVE/DEAD stain (Thermo Fisher Scientific, Waltham, MA), lifted in Accutase at 37°C for 30 minutes, and run through an Attune NxT Flow Cytometer (Thermo Fisher Scientific, Waltham, MA). Cells were treated with the Foxp3/Transcription Factor Staining Buffer Set (Thermo Fisher Scientific, Waltham, MA, USA) according to manufacturer’s recommendations and stained for 1 hour at 4°C with the following antibodies: NGFR (345110; 1:600, BioLegend, San Diego, CA, USA), α-tubulin (NB100-69AF405, 1:300, Novus, Centennial, CO, USA) and CD66c (12-0667-42, 1:600, Invitrogen, Waltham, MA). GFP expression was gated on live cells.

### Immunofluorescence staining

Human airway epithelial cultures, NHP explants, or murine lung sections were immunostained for ciliated cells using acetylated α-tubulin (1:200, D20G3, Cell Signaling Technology, Danvers, MA, USA), or secretory cells using SCGB1A1 (aka CCSP, 1:100, 07-623, EMD Millipore). Primary antibodies were detected using a goat anti-rabbit Alexa 546 secondary antibody (1:600, A-11035, Thermo Fisher Scientific, Waltham, MA, USA). Alexa Fluor 647 Phalloidin was used to stain F-actin (1:50, A22287, Invitrogen).

### LDH release assay

LDH release was quantified according to the manufacturer’s recommendations (LDH-Glo Cytotoxicity Assay, Promega, Madison, WI). Briefly, basolateral media from each condition was collected in a 96 well plate. The LDH detection reagents were mixed in a 1:1 ratio and applied to each well and incubated for 30 minutes. Luminescence was recorded using a SpectraMax i3x plate reader. Negative control was an untreated HAE and Max control was included as described in the LDH-Glo Cytotoxicity kit.

### Non-human primate explants

Non-human primate tracheas were collected at the Children’s Hospital of Philadelphia. Explants were cultured on Surgifoam and maintained in Pneumacult ALI-M media. Explants were transduced with various AAV capsids (2x10^10^ vg) formulated in 5% NaCl by inverting the tissue on plastic for 2 hours as previously described^20^. Explants were left untreated or treated with 1 μM Dox 48 hours post-transduction. GFP expression was visualized 2 weeks post-transduction.

### Delivery to murine airways

6-8 week old BALB/c mice were anesthetized using 2% isofluorane and up to 100 µl of 10^11^ vg of each capsid formulated in 6% NaCl was instilled intratracheally. Mice were humanely euthanized 2 weeks later and lungs were inflated with 4% paraformaldehyde, passed through a sucrose gradient, and embedded for sectioning. Tissues sections were stained for club cells (SCGB1A1) and ciliated cells (tubulin).

## Acknowledgements

We thank the University of Iowa Cell and Tissue Culture Core for providing cultured airway epithelial cells for this project. This work was supported by Emily’s Entourage (McCray and Davidson), the Boomer Esiason Foundation (Cooney), the Cystic Fibrosis Foundation (MCCRAY25G0), and NIH P01HL152960.

---

**Supplemental Figure 1:**
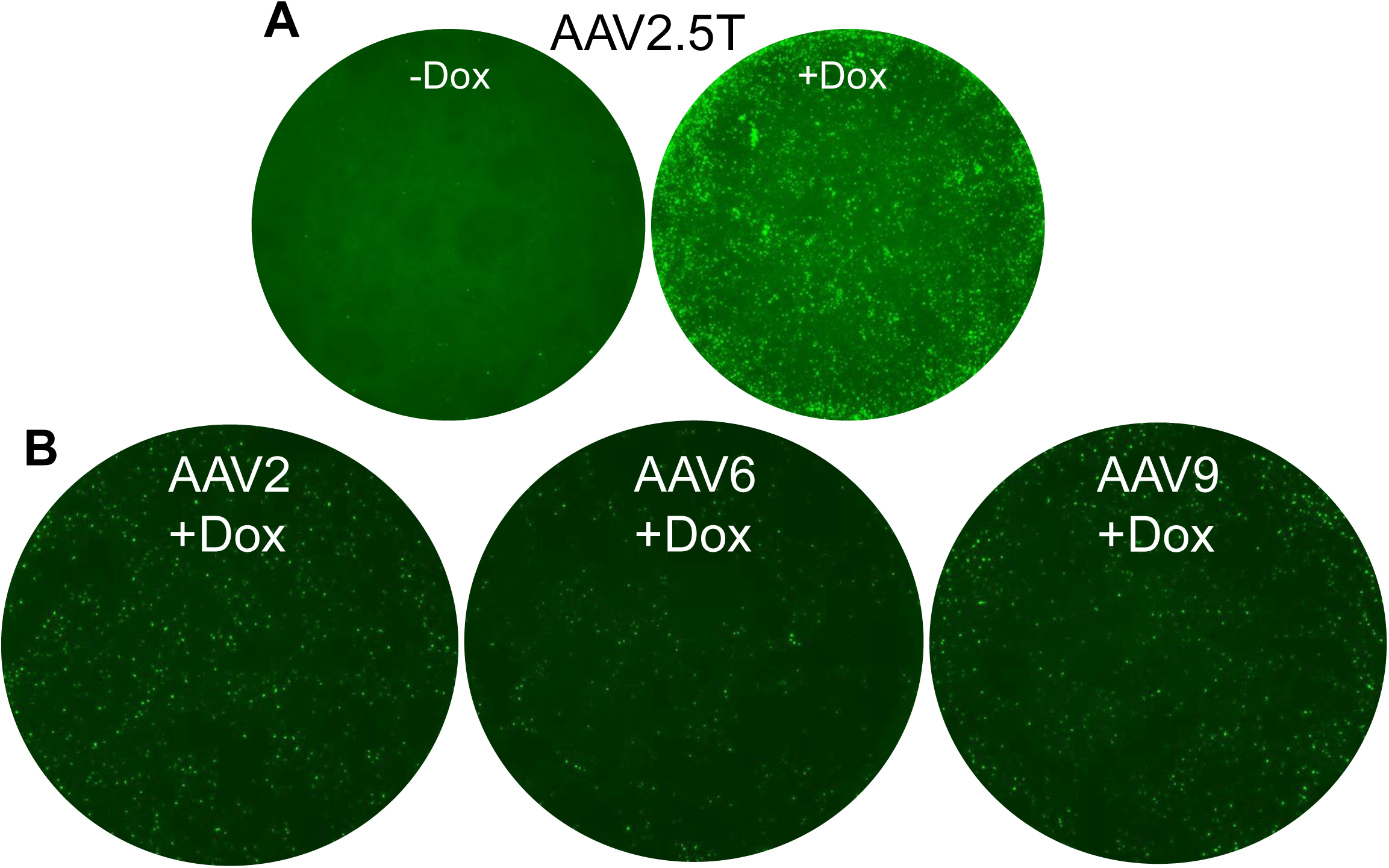
GFP expression in HAE cultures for AAV2.5T and parental serotypes. A) AAV2.5T-GFP (Spirovant) was tested on HAE overnight (MOI=50,000) with and without Dox (1 µM) 48 hours later. Representative images from airway cultures imaged 2 weeks later. B) Parental serotypes were tested on HAE overnight (MOI=50,000) and with Dox (1 µM) 48 hours later. Representative images from airway cultures imaged 2 weeks later.

**Supplemental Figure 2:**
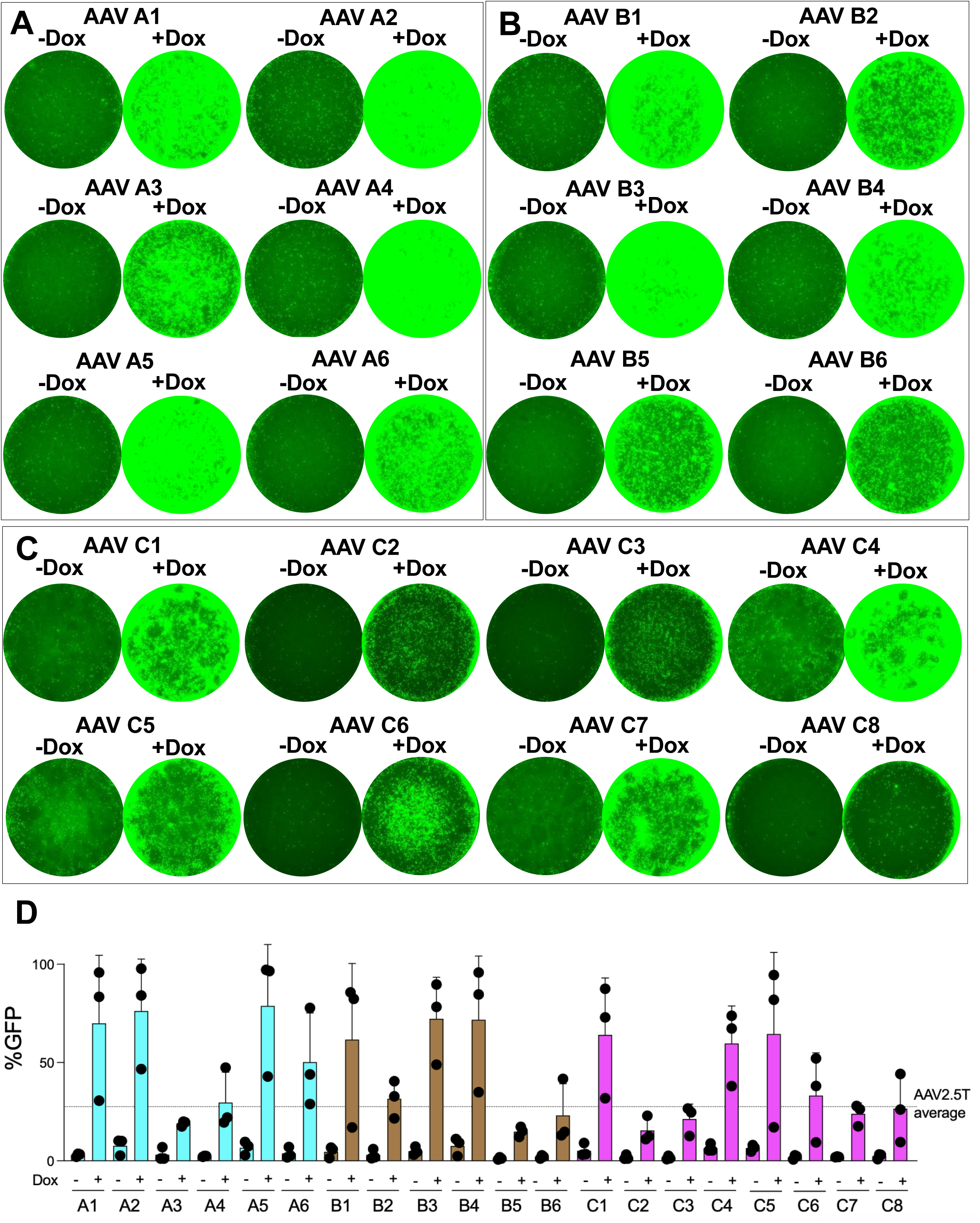
Transduction properties of the AAV variants on human airway epithelia (HAE). A-C) HAE were transduced with A1-A6, B1-B6, and C1-8 (MOI=50,000) overnight. 2 days post-transduction, cells with the “+Dox” designation were treated overnight with 1 µM doxorubicin (Dox). Images were acquired 2 weeks post-transduction. D) Flow cytometry quantification of %GFP in each culture, n=3. Dotted line indicates AAV2.5T average from Figure 2A, used as a benchmark for HAE transduction and to help identify the top 10 AAV-AE variants.

**Supplemental Figure 3:**
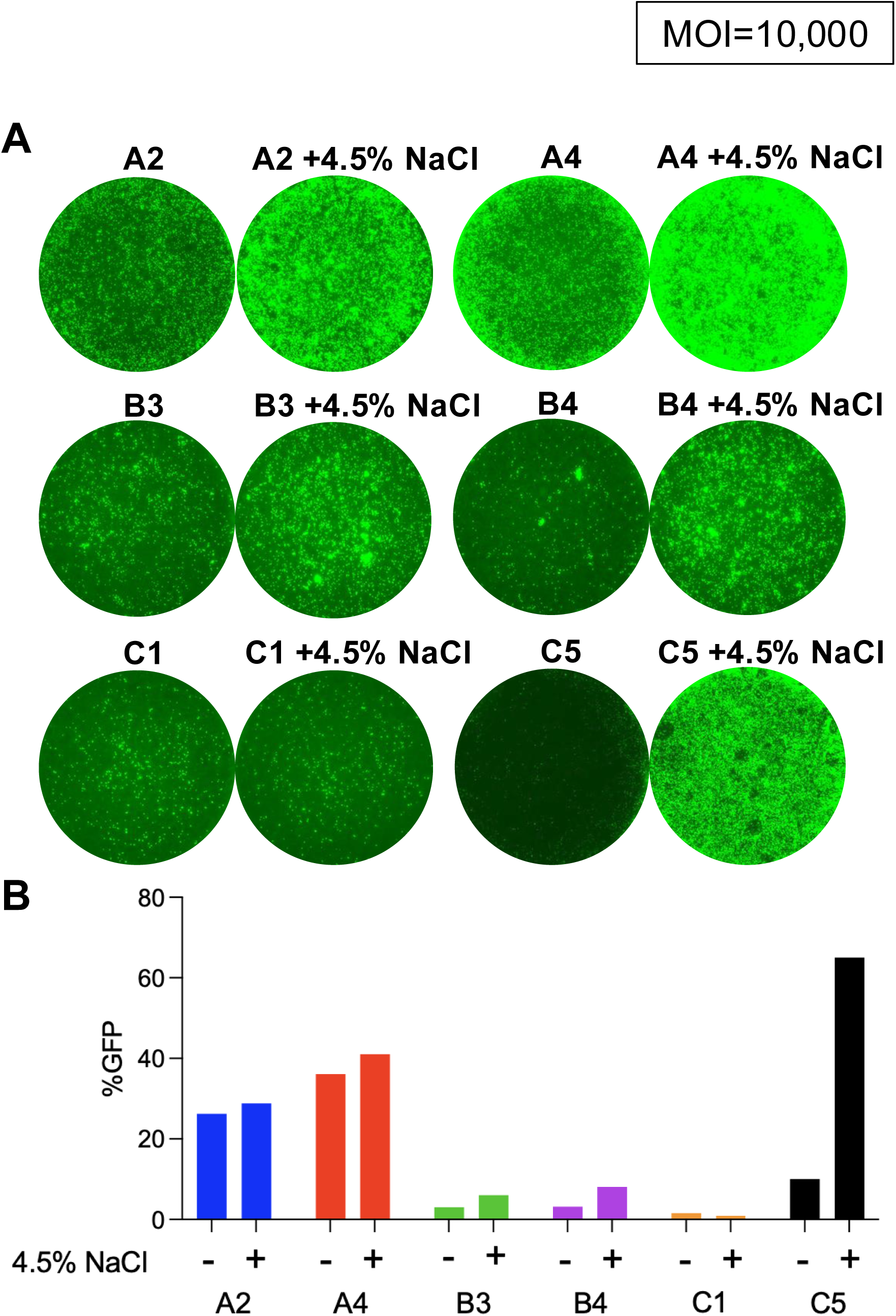
Formulating AAVs in hypertonic saline further increases HAE transduction of various capsids. AAV capsids A2, A4, B3, B4, C1 and C5 (MOI=10,000) were tested with and without 4.5% NaCl (2 hour apical transduction) and Dox (1 µM, 48 hours post-transduction). A) GFP was visualized by fluorescence microscopy 2 weeks post-transduction and B) quantified by flow cytometry (n=1).

**Supplemental Figure 4:**
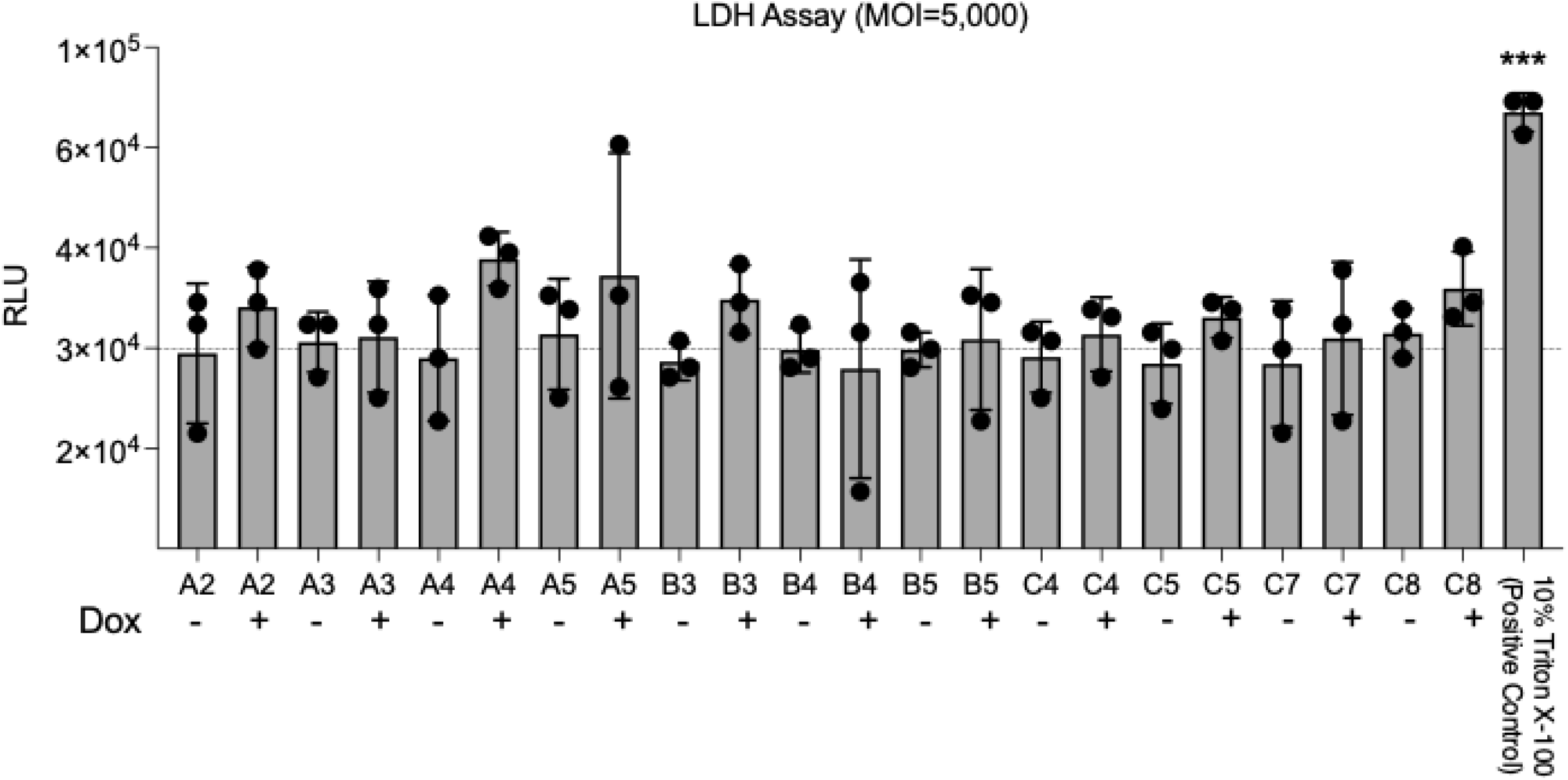
GFP does not cause cytotoxicity. AAV capsids were transduced at an MOI=5,000 and treated with or without 1 μM Dox. 2 weeks post-transduction, an LDH assay was performed to quantify levels of cytotoxicity. Luminescence was measured and is reported at relative light units (RLU). Dotted line indicates untreated control, n=3, ***p<0.0005.

**Supplemental Table 1:** Summary of AAV capsids and titers.

| Capsid | Vector name | Titer (vg/ml) |
| --- | --- | --- |
| A1 | AAV-AE-A1 iCAG.eGFP | 2.83E+11 |
| A2 | AAV-AE-A2 iCAG.eGFP | 7.56E+11 |
| A3 | AAV-AE-A3 iCAG.eGFP | 1.58E+12 |
| A4 | AAV-AE-A4 iCAG.eGFP | 1.14E+12 |
| A5 | AAV-AE-A5 iCAG.eGFP | 5.79E+11 |
| A6 | AAV-AE-A6iCAG.eGFP | 3.63E+11 |
| B1 | AAV-AE-B1 iCAG.eGFP | 7.83E+11 |
| B2 | AAV-AE-B2 iCAG.eGFP | 1.55E+12 |
| B3 | AAV-AE-B3 iCAG.eGFP | 1.03E+12 |
| B4 | AAV-AE-B4 iCAG.eGFP | 1.03E+12 |
| B5 | AAV-AE-B5 iCAG.eGFP | 1.89E+12 |
| B6 | AAV-AE-B6 iCAG.eGFP | 8.83E+11 |
| C1 | AAV-AE-C1 iCAG.eGFP | 6.96E+12 |
| C2 | AAV-AE-C2 iCAG.eGFP | 9.74E+12 |
| C3 | AAV-AE-C3 iCAG.eGFP | 8.67E+12 |
| C4 | AAV-AE-C4 iCAG.eGFP | 8.16E+12 |
| C5 | AAV-AE-C5 iCAG.eGFP | 3.48E+12 |
| C6 | AAV-AE-C6 iCAG.eGFP | 3.00E+12 |
| C7 | AAV-AE-C7 iCAG.eGFP | 5.40E+12 |
| C8 | AAV-AE-C8 iCAG.eGFP | 4.07E+12 |

